# A multivariate phylogenetic comparative method incorporating a flexible function between discrete and continuous traits

**DOI:** 10.1101/776617

**Authors:** Yuki Haba, Nobuyuki Kutsukake

## Abstract

One major challenge of using the phylogenetic comparative method (PCM) is the analysis of the evolution of interrelated continuous and discrete traits in a single multivariate statistical framework. In addition, more intricate parameters such as branch-specific directional selection have rarely been integrated into such multivariate PCM frameworks. Here, originally motivated to analyze the complex evolutionary trajectories of group size (continuous variable) and social systems (discrete variable) in African subterranean rodents, we develop a flexible approach using approximate Bayesian computation (ABC). Specifically, our multivariate ABC-PCM method allows the user to flexibly model an underlying latent evolutionary function between continuous and discrete traits. The ABC-PCM also simultaneously incorporates complex evolutionary parameters such as branch-specific selection. This study highlights the flexibility of ABC-PCMs in analyzing the evolution of phenotypic traits interrelated in a complex manner.

## Introduction

Phylogenetic comparative methods (PCMs) provide a powerful statistical framework for investigating the patterns and processes of trait evolution (Felsenstein 1985; Harvey & Pagel 1991; Nunn 2011; Garamszegi 2014). The recent development of PCMs permits analyses of biologically interrelated discrete and continuous variables in a single multivariate statistical framework (Table 1). The development of such multivariate PCMs is crucial for two reasons. First, conducting two separate univariate analyses for a continuous trait and a discrete trait is redundant when the two variables are interrelated. Second, and more importantly, separate univariate analyses will miss the opportunity to consider important biological links between these traits.

**Table 1.** Summary of previous methods and the originality of this study.

| Trait | Approach |  |
| --- | --- | --- |
|  | Univariate models | Multivariate models (link function) |
| Discrete | Mk model<sup>1</sup> | Mk model<sup>1</sup> |
|  | Threshold model<sup>2*</sup> | Threshold model<sup>2*</sup> |
|  | PLogReg<sup>3</sup> (w/o independent variables) |  |
| Continuous | PLogReg<sup>3</sup> | MCMCglmm<sup>4,5</sup> |
|  | MCMCglmm<sup>4,5</sup> |  |
| Both discrete and continuous |  | Threshold model<sup>2</sup><br>(determined by a threshold)* |
|  | - | PLogReg<sup>3</sup> (logit link) |
|  |  | MCMCglmm<sup>4,5</sup> (logit and probit link) |
|  |  | This study<br>(researcher-defined function) |

<sup>1</sup>Pagel 1994; Lewis 2001
<sup>2</sup>Felsenstein 2012
<sup>3</sup>Phylogenetic logistic regression, Ives and Garland 2010
<sup>4</sup>Hadfield and Nakagawa 2009
<sup>5</sup>Hadfield 2015
\*continuous trait (liability) is unobservable

The threshold model (Felsenstein 2005, 2012) was among the first PCMs to fully combine both discrete and continuous traits. The idea of the threshold model was originally developed in quantitative genetics by Wright (1934) to understand how multiple underlying genetic loci contribute to categorical traits such as the number of digits in guinea pigs. The threshold model assumes an unobservable continuous trait called ‘liability’ that underlies a discrete trait of interest. Because the liability is a continuous trait, Brownian motion has been conventionally used to model its evolution. Then, the state of the discrete trait of interest is determined by whether the liability trait value is below or above a particular threshold. This model allows users to incorporate both continuous and discrete traits in a straightforward way, as well as to estimate the covariance between the liability trait and other continuous traits of interest. However, because liability is unobservable, it is impossible to directly infer a latent function between the discrete trait and other observable continuous traits of interest. Moreover, although it is convenient to assume that the discrete trait is determined by a simple threshold (which can be treated as a probit link function in a framework of a phylogenetic generalized linear mixed model, PGLMM; Hadfield 2015; also see Ives and Garland 2014), it is unclear if the assumption is always biologically valid. At the very least, it is desirable for researchers to be able to assume other forms of latent functions (Fig. 1).

**Fig. 1.**
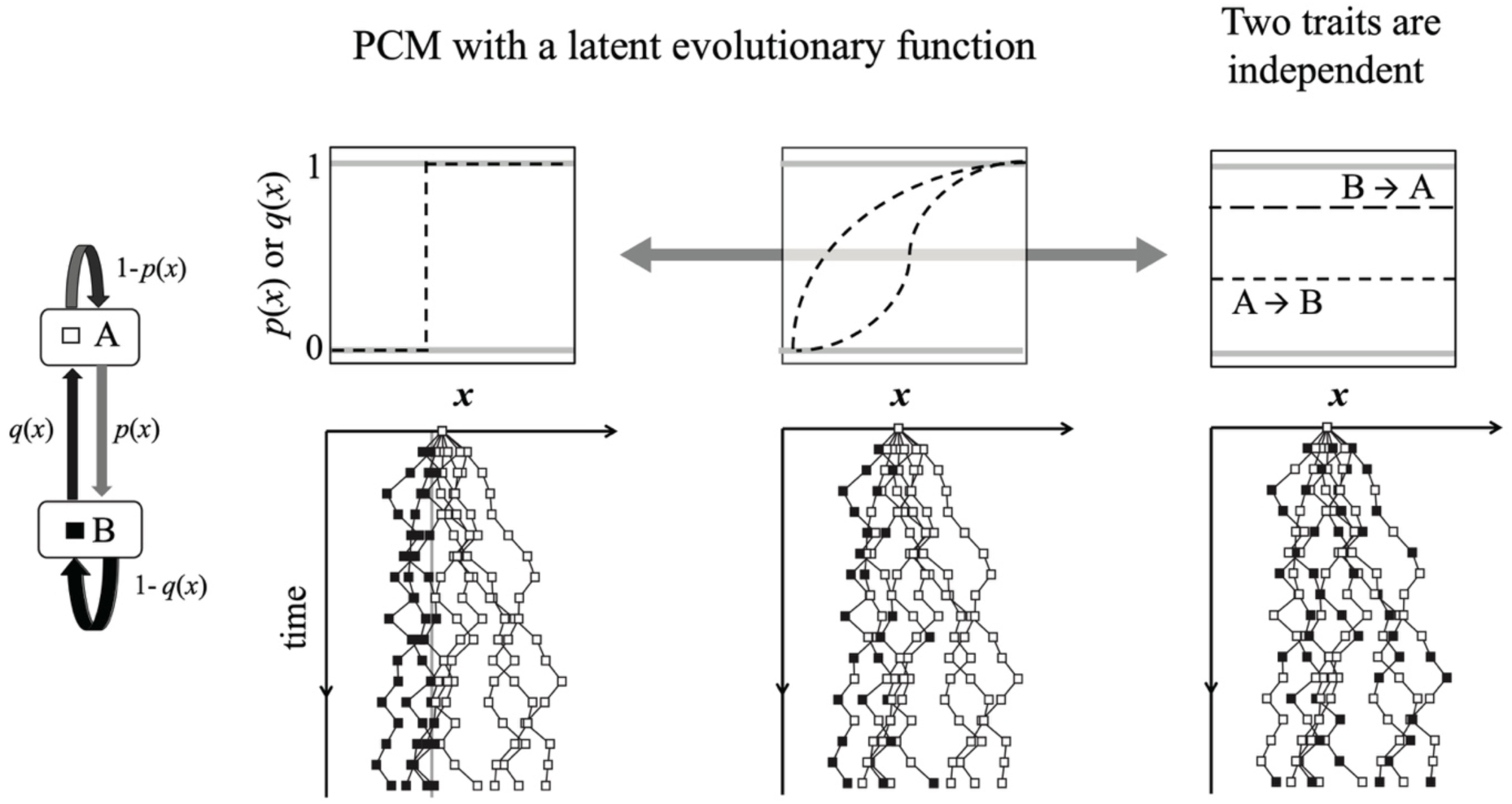
Variation in multivariate PCMs with a latent function between discrete and continuous traits. Conventional PCMs can explore various latent evolutionary functions between continuous and discrete traits (Table 1). Here, we consider a simple example of the evolutionary link between a continuous and a discrete trait. The continuous trait (*x*) evolves by a process similar to Brownian motion. The discrete trait, on the other hand, evolves according to the transition probabilities *p*(*x*) and *q*(*x*), i.e., the latent evolutionary functions. The state of the discrete trait at each time step is shown as a white (A) or black (B) square. At one extreme, the transition of discrete states is determined by a certain “threshold” value of the continuous trait. At the other extreme, the discrete trait and the continuous trait are independent. In such a case, the transition probabilities do not vary with the continuous traits (right). In our framework, any functions between the two extreme cases can be incorporated based on biological hypotheses (middle). Furthermore, *x* can be either a measurable trait or a latent value. Here, logistic and exponential functions are shown as examples.

In addition to Felsenstein’s threshold model, other methods that can link discrete traits and continuous traits have been proposed (Ives & Garland 2010, 2014; Hadfield and Nakagawa 2009; Hadfield 2015; see Table 1). For example, Ives and Garland (2010) developed a phylogenetic logistic regression to test the effects of observable continuous independent traits on a discrete dependent trait (phylogenetic logistic regression, Ives & Garland 2010). Hadfield and Nakagawa proposed a phylogenetic generalized linear mixed model employing a Bayesian approach (MCMCglmm, Hadfield and Nakagawa 2009; Hadfield 2015; also see Ives and Garland 2014 for other approaches and a comparison of their performance). These approaches successfully extended traditional linear models to enable nonlinear link functions (logit function in a phylogenetic logistic regression; logit or probit functions in MCMCglmm) between discrete and continuous traits. Notably, a probit-GLM can be mathematically equivalent to the threshold model (see Hadfield 2015). However, limitations to these models still exist. For example, the model by Ives and Garland (2010) assumes that the continuous traits are known values measured empirically and does not allow the continuous traits to evolve along the tree (see Felsenstein 2012 and Hadfield 2015 for models that do not have such assumption; but also see Ives and Garland 2014 for analyses of the relatively small effects of phylogenetic signal on continuous traits). Moreover, the link function in the models is predetermined. In most cases, equipped functions (e.g., logit or probit functions) are those that have useful mathematical properties for analyzing a relationship between discrete and continuous traits. Still, cases may exist in which it is biologically valid to establish a more complicated function between discrete and continuous traits. Therefore, it is ideal to prepare a framework that enables researchers to examine a flexible function to test specific hypotheses of interest.

Furthermore, analyses will be even more complex if additional variables of interest are included, such as the presence of branch-specific directional selection (Kutsukake and Innan 2013, 2014). This complication often prevents the description of a likelihood function of the model, which most conventional PCMs require (e.g., the maximum likelihood, Bayesian approach). The aforementioned PCMs (Felsenstein 2005, 2012; Ives and Garland 2010, 2014; Hadfield and Nakagawa 2009; Hadfield 2015) cannot incorporate branch-specific directional selection into their models.

Here, we propose a PCM using approximate Bayesian computation (ABC) to overcome the difficulties discussed above (Fig. 2). ABCs have been shown to facilitate flexible analyses in a comparative framework and therefore have increasingly been applied to PCMs with intricate evolutionary scenarios (Bokma 2010; Slater *et al*., 2012; Kutsukake and Innan 2013, 2014; Janzen *et al*. 2015; Harano & Kutsukake 2018). Briefly, an ABC-PCM estimates parameters of interest by simulating phenotypic evolution without a likelihood function (Beaumont 2010; Csillery *et al*. 2010). The proposed parameters are accepted only when the simulated data and real data are similar, and the accepted data comprise posterior distributions of parameters. Thanks to this flexibility, ABC-PCMs enable researchers to test evolutionary models whose likelihood function is mathematically intractable.

**Fig. 2.**
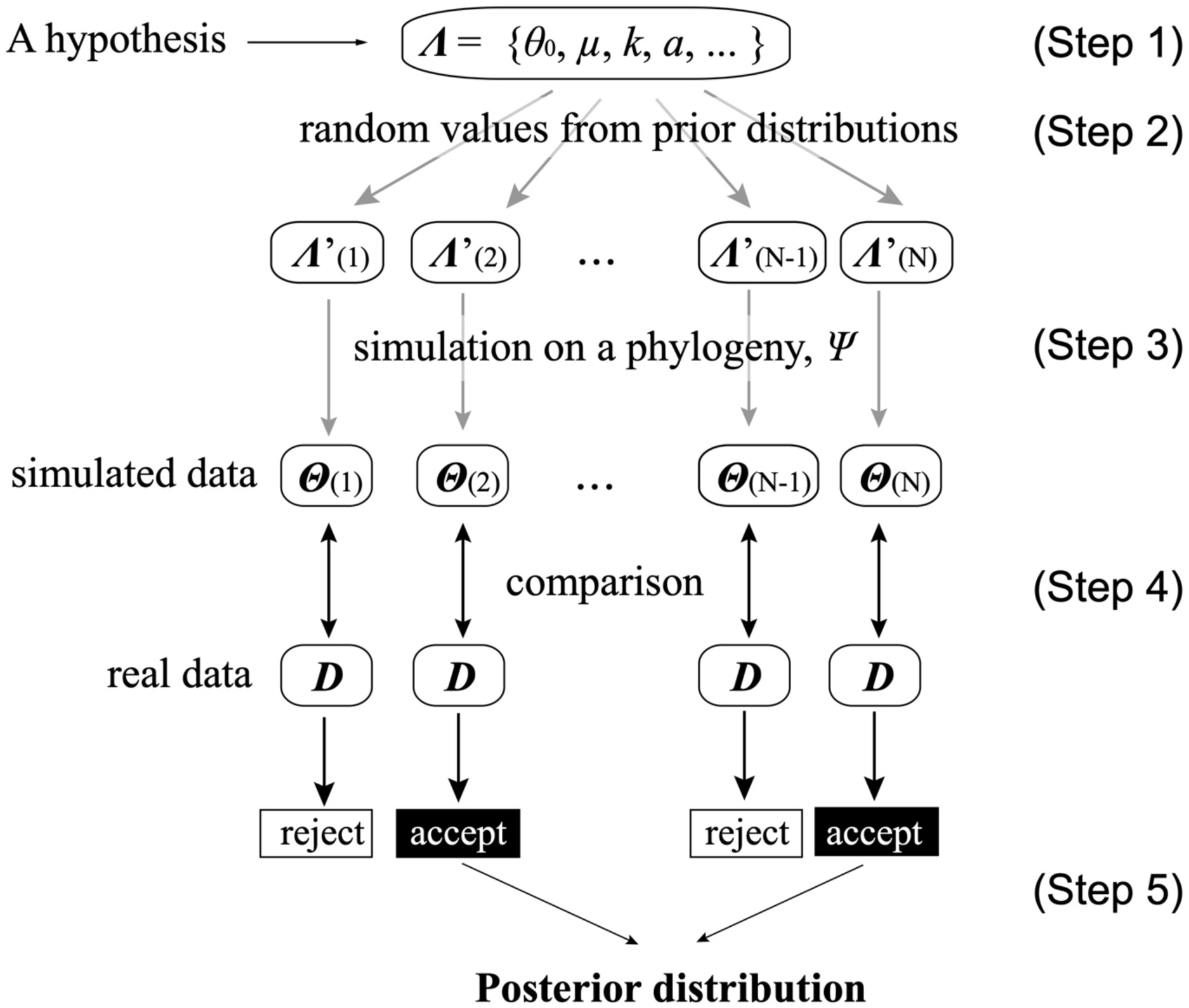
Schematic of our ABC-PCM approach. First, determine parameters of interest and set prior distributions (Step 1). Example parameters include phenotype of the MRCA (*θ*_0_), evolutionary rate (*μ*), directional selection parameter (*k*), latent function parameter (*a*), and so on. Next, generate random values for parameters from the prior distributions (Step 2). Trait simulations on a phylogeny *Ψ* are then conducted (Step 3), and the simulated data ***Θ*** are compared to real data ***D*** to determine whether the data are accepted or rejected (Step 4). A number (N) of simulations are conducted until enough samples are collected to infer the posterior distribution (Step 5).

Our initial motivation for extending ABC-PCMs was to analyze a heterogeneous evolutionary pattern between group size and social system in African subterranean mole rats (family Bathyergidae; Table 2). In this clade, species have varied social structures that span solitary, social, and eusocial states. Eusociality has independently evolved twice: in naked mole rats (*Heterocephalus glaber*) and in Damaraland mole rats (*Fukomys damarensis*) (Jarvis 1981; Sherman *et al*. 1991; Bennett & Faulkes 2000; Faulkes & Bennett 2013). We hypothesized that the probabilities of evolutionary transitions among eu/social and solitary states (discrete trait) change depending on group size (continuous trait). To capture all of these complex evolutionary features, we are required to incorporate (1) the evolutionary trajectory of sociality and group size, (2) the parameters for the trait-dependent latent functions, and (3) the presence of branch-specific directional selection on group size in the two eusocial species. Certain aspects of these features can be analyzed using previous methods; however, it is not possible to incorporate all factors in a single model with those methods. Then, based on the example data, we discuss how this ABC-PCM can be used for inferring other similar complex evolutionary processes.

**Table 2.** Data used in this study.

| Species | Social system | Group size, mean (SD) |
| --- | --- | --- |
| Naked mole rat, *Heterocephalus glaber*<sup>1</sup> | eusocial | 75 (48.65)<sup>A</sup> |
| Damaraland mole rat, *Fukomys damarensis*<sup>1</sup> | eusocial | 11 (6.26) |
| Whyte's mole rat, *Fukomys hottentotus*<sup>1</sup> | social | 5.16 (2.62)<sup>B</sup> |
| Mechow's mole rat, *Fukomys mechowii*<sup>2</sup> | social | 9.91 (2.49) |
| Mashona mole rat, *Fukomys darlingi*<sup>3</sup> | social | 7 (not available<sup>C</sup>) |
| Ansell's mole rat, *Fukomys ansellii*<sup>4</sup> | social | 8.72 (2.15) |
| Cape mole rat, *Georychus capensis*<sup>5</sup> | solitary | 1<sup>D</sup> |
| Cape dune mole rat, *Bathyergus suillus*<sup>5</sup> | solitary | 1<sup>D</sup> |
| Namaqua dune mole rat, *Bathyergus janetta*<sup>5</sup> | solitary | 1<sup>D</sup> |
| Silvery mole rat, *Heliophobius argenteocinereus*<sup>5</sup> | solitary | 1<sup>D</sup> |

<sup>1</sup>Bennett & Faulkes 2000; <sup>2</sup>Sichilima *et al.* 2008; <sup>3</sup>Bennett *et al.* 1994; <sup>4</sup>Sichilima *et al.* 2011; <sup>5</sup>Burda & Kawalika 1993; see Van Daele *et al.* (2013) for a recent classification.
<sup>A</sup>Because group sizes were categorized at an interval of 25 individuals, we used the mean value of each range for calculating the mean and SD (e.g., 37.5 for groups of 25 to 50 individuals).
<sup>B</sup>When calculating the mean group size of this species, we used a group size of 11 when it exceeded 10, because the original data for group size pooled group sizes larger than 10 into ">10" (p. 92 in Bennett & Faulkes 2000).
<sup>C</sup>Because no data on intraspecific variation were available, we used the mean value of the SD, 2.42, of the other three social species.
<sup>D</sup>The mean litter size ranges from 2.46 to 5.94 in solitary species (Jones *et al.* 2009). To account for this, we set a value of 2.56 as their SD to cover the range of possible group sizes.

## Methods

### ABC-PCM

Our ABC-PCM extends a previously developed framework (Fig. 2; Kutsukake & Innan 2013, 2014) designed to analyze heterogeneous evolutionary modes whose likelihood is not straightforward to describe. We assume knowledge of the species tree *Ψ*, which consists of information on tree shape and the lengths of all branches in the topology. We also assume that trait data at the tips of the tree ***D*** have been observed.

In bivariate analyses of continuous and discrete traits, several causational patterns are possible. For example, a discrete trait can be determined by a continuous trait that evolves on its own, or vice versa. Alternatively, it is also possible to assume no *a priori* causality between the continuous or discrete traits (Hadfield and Nakagawa 2009; Ives and Garland 2010). Any of these cases can be modeled by the ABC-PCM framework proposed in this paper. Moreover, this framework can be used regardless of whether a given value is latent (e.g., liability in the threshold model) or measurable.

We hereafter consider a simple case of interrelated evolution in which a continuous trait determines a discrete trait. In our example study of African subterranean mole rats, both traits were measurable.

Briefly, our ABC-PCM process is implemented as follows (see Fig. 2 for a visual schematic).

Let ***Λ*** be the parameter set to be estimated based on a hypothesis.

(Step 1) Determine prior distributions for all parameters in ***Λ***. If prior biological knowledge is available, it can be used to set informative (i.e., strong) prior distributions. (Step 2) Parameters used in the simulation (***Λ*’**) are randomly generated from the prior distributions.

(Step 3) Using ***Λ*’**, the trait evolution is simulated on the phylogeny *Ψ*. In the current model, the trait simulation has two parts corresponding to continuous and discrete trait evolution (Fig. 1).

(Step 3a) The continuous trait evolves via Brownian motion (Felsenstein 1985; note that other evolutionary models can be used; see Kutsukake and Innan 2013).

(Step 3b) Then the discrete trait is determined according to the probability of transition between two states (*A* and *B*) as a function of the continuous trait. Note again that this setting can be relaxed such that causation between continuous and discrete traits can be flexibly changed according to a hypothesis of interest. Here, *p*(*x*) is the probability function that state *A* changes to *B* given the continuous trait *x*. Similarly, *q*(*x*) is the probability that state *B* changes to *A* given the continuous trait *x* (Fig. 1).

(Step 4) Calculate the likelihood by comparing simulated data ***Θ*** with the real data ***D*** and determine whether the parameter set is accepted or rejected. A joint probability (full likelihood) for the comparison of *n* species can be calculated. In most cases, this probability is difficult to obtain. In such cases, a composite likelihood that is proportional to the full likelihood can be used as an approximate proxy. Intraspecific variation in the trait data can be considered by assuming a certain distribution for the trait when calculating the likelihood.

Then the acceptance or rejection of the parameters can be determined based on the likelihood. Several methods of judgment exist (Marjoram *et al*. 2003; Marjoram and Tavare 2006).

(Step 5) Repeat Steps 2–4 until a sufficient number of parameters ***Λ*’** is accepted. Then, posterior distributions and credible intervals can be estimated.

Although the fundamental structure of ABC is straightforward, the number/choice of summary statistics and the width of tolerance for judging the acceptance or rejection of simulated data at Step 4 are controversial, and there is no general consensus on the choice of summary statistics and tolerance (Beaumont *et al*. 2002; Csillery *et al*. 2010; Leuenberger & Wegmann 2010). In this study, we use a combination of perfect match and joint probability as summary statistics (see *Acceptance and summary statistics* below for details).

### Application—social evolution in African mole rats

One example of complex evolution where continuous and discrete traits are interrelated is social evolution. Sociality in animals can be characterized using a discrete classification based on mating and/or social systems (e.g., Shultz *et al*. 2011). Average group size of a species, a continuous trait, is also an important variable in characterizing sociality in animals (e.g., comparative analyses: Faulkes *et al*. 1997; Sheehan *et al*. 2015). Loss of sociality is correlated with a decrease in group size (secondary loss of sociality; Wcislo & Danforth 1997; Beauchamp 1999; Sheehan *et al*. 2015). Likewise, cooperative social systems are likely to appear as the group size increases (e.g., limited dispersal by ecological constraints: Emlen 1982; Duffy & Macdonald 2010). Thus, the complex social evolution across African mole rats is an ideal system to test our framework. Here, species-specific sociality and the mean group size for each species (*x*) are the target traits.

### Dataset

We surveyed the literature for field data on the sociality (solitary, social, or eusocial), mean group size, and/or distribution of group sizes in each species (Table 2). When the distribution of group size was available, we calculated its mean and standard deviation for each species. For solitary species, we regarded the mean group size as one. In reality, however, their group sizes can deviate from this value, because females may have dependent, pre-dispersal offspring; thus, the group size of solitary species can also have a distribution. Therefore, we incorporated a realistic value for the variance of solitary species (Table 2). We used the phylogeny presented in Faulkes *et al*. (2004), who used mitochondrial genes 12s rRNA and cyt *b*. The mean value of the estimated divergent interval in millions of years was used as the length of each branch.

### Parameter set and trait simulation

The trait simulation included six parameters to be estimated: the group size of the most recent common ancestor (MRCA) (*θ*), the rate of evolution (*µ*), the parameters of directional selection (*k*_*n*_ and *k*_*d*_), and the parameters for latent functions (*a* and *b*, or *c* and *d*, depending on the model used; see next section). Table 3 shows the notation and settings of the prior distributions.

**Table 3.** Parameters estimated in this study.

| Parameter | Notatio<br>n | Prior<br>distribution | Posterior distribution (95% CI) |  |
| --- | --- | --- | --- | --- |
|  |  |  | Model 1<br>(logistic) | Model 2<br>(exponential) |
| Most recent common ancestor (MRCA) | $\theta$ | U(1.001, 10) | 1.1–5.22 | 2.45–9.15 |
| Baseline evolutionary rate (per million years) | $\mu$ | U(0.001, 30) | 6.4–29.5 | 3.62–28.3 |
| Directional selection (naked mole rats) | $k_n$ | U(0.001, 30) | 0.99–3.57 | 1.01–1.93 |
| Directional selection (Damaraland mole rats) | $k_d$ | U(0.001, 5) | 0.82–4.22 | 0.82–4.64 |
| Curvature of transition functions (solitary ⇌ social) | $a$ or $c$ | U(-15,15) | $a$: 1.52–14.50 | $c$: 1.28–14.60 |
| Position of transition functions | $b$ or $d$ | U(0,10) | $b$: 1.52–4.94 | $d$: 1.28–5.07 |

The evolutionary process of the continuous trait was as follows. First, we generated the values of six parameters from the prior distributions. Then, based on the proposed value of *x* at the root (i.e., the group size of the MRCA, *θ*), we randomly determined the state (i.e., sociality) of the MRCA by either *p*(*x*) or *q*(*x*). The number of evolutionary events that change traits (i.e., either an increase or decrease in group size) on each branch was respectively modeled as a random number from a Poisson distribution with a mean equaling the product of the evolutionary rate, *µ* (trait change per million years), and branch length *τ* (millions of years). The degree of trait change by one evolutionary event is a random value from an exponential distribution, with its mean being an arbitrary value *φ* (set to 0.01). *φ* corresponds to the average effect of one change on the trait value. We used an exponential distribution because a change in trait due to one evolutionary event would have small effects on traits in most cases, and large effects less frequently. Thus,

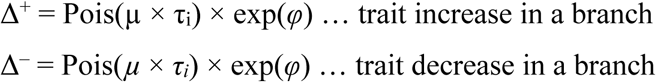

where *τ*_*i*_ is the length of the *i*th branch. The total trait change on the branch is calculated as Δ^+^ – Δ^−^.

Changes in group size caused by one evolutionary event would depend on the initial group size. For example, an increase in the group size by one individual will have different biological meanings in solitary and eusocial species. Therefore, we transformed the value of group size to a natural logarithmic scale during trait simulation. Because the mean group size *x* cannot be less than one, we enforced a lower bound of one. That is, when a trait change resulted in a value less than one on a branch, the change was not implemented.

We also tested whether there were selective pressures for increasing group size in the two branches leading to eusocial species. In our ABC-PCM model, the parameter *k* represents directional selection (Kutsukake & Innan 2013, 2014). The selection parameter *k* biases the number of positive or negative trait changes such that the number of trait changes Pois(*µ* × *τ*_*i*_) × exp(*φ*) is multiplied or divided by *k*, respectively (see Kutsukake & Innan 2013 for more details). Thus,

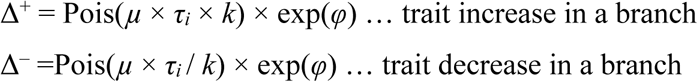

and again, the total trait change on the branch is calculated as Δ^+^ – Δ^−^.

When *k* = 1, the evolutionary mode of trait evolution is asymptotically identical to the Brownian motion. On the other hand, a significant departure of *k* from 1 is used as a signature of directional selection.

In this analysis, we set selection parameters for the branches for both naked mole rats and Damaraland mole rats as *k*_*n*_ and *k*_*d*_, respectively (Table 3), to test our hypothesis that branch-specific selective pressure has increased group size in eusocial species (Jarvis & Bennett 1993; Young *et al*. 2015). When the 95% CI of the posterior distribution of *k* was larger than 1, it was considered a signature of directional selection for larger group size.

### Models of latent evolutionary functions

At each evolutionary step, the discrete trait value was determined by the continuous trait value according to the latent functions (see Step 3b of *ABC-PCM*). Because the shape of this function between sociality and group size was unknown, we took advantage of the flexibility of our ABC-PCM framework and tested two examples of evolutionary models with different transition functions, *p*(*x*) and *q*(*x*). We note that this framework is not limited to these two models but can use any function, depending on the focal biological traits and hypotheses of interest. For simplicity, we set the latent evolutionary functions such that *p*(*x*) + *q*(*x*) = 1, but this is not required if one desires otherwise.

### *Model 1:* Logistic function

In this model, we used a logistic function as the latent evolutionary function, which was similar to a previous model using a generalized linear model for a binary dependent term (Ives and Garland 2010). The transition functions *p*(*x*) and *q*(*x*) were defined as follows:

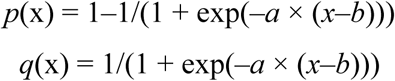

The parameter *a* determined the curvature of *p*(*x*) and *q*(*x*), i.e., the effects of the group size on sociality (Fig. 1). When *a* is equal to 0, *p*(*x*) and *q*(*x*) are flat and the transition between social and solitary is independent of group size (Fig. 1, right). When *a* is positive, a species with large group size is likely to be social; when *a* is negative, a species with large group size is likely to be solitary. Importantly, when |*a*| is large enough, the transition between social and solitary is determined by a certain group size, which virtually behaves like a step function (Fig. 1, left). The parameter *b* is the *x* value at which *p*(*x*) and *q*(*x*) equal 0.5, and determines the group size at which the probability of transitioning from solitary to social becomes larger than the probability of the reverse. Because the curvature and midpoint of *p*(*x*) and *q*(*x*) (i.e., *a* and *b*) were unknown *a priori*, we set broad prior distributions of *a* and *b* (Table 3).

### *Model 2:* Exponential function

In the second model, we used an exponential function as the latent evolutionary function. We set the transition functions *p*(*x*) and *q*(*x*) as follows:

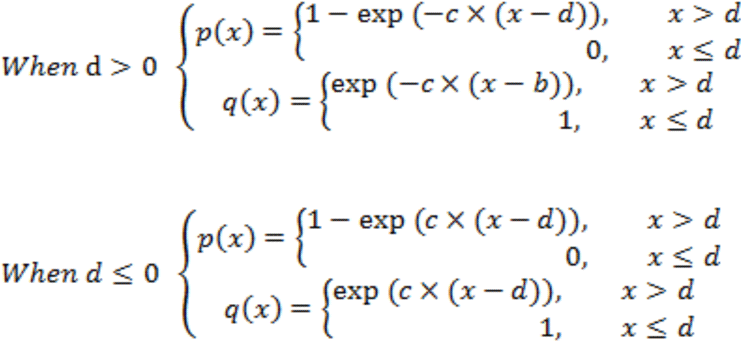

These functions describe an exponential decrease or diminishing increase in the transition probability, and a given species with a mean group size of one is always a solitary species. Similar to *a* and *b* in model 1, *c* and *d* determine the curvature of the exponential function and the point at which the exponential decrease/increase begins, respectively. Note that one side of the exponential function used here is asymptotically equal to either 0 or 1 when *x* is sufficiently large, but that of the other side is not; therefore, we set the probability of state change as 0 or 1 when *x* is smaller than *d*, at which the functions reach 0 or 1, respectively.

### Acceptance and summary statistics

Among ABC-PCMs, no clear consensus has been reached concerning the number/choice of summary statistics or the width of tolerance for judging the acceptance or rejection of simulated data. In the present analyses, we used two summary statistics to assess the fit of the simulated data to actual data. First, for the discrete variable (i.e., sociality), we only accepted simulations in which the simulated data were a perfect match to the real data. For the continuous variable, we used a conventional method that uses a joint probability (Kutsukake & Innan 2013, 2014); we first calculated the probability that the real trait value is gained under a simulated trait value for all 10 species. When calculating the probability, group size (back-transformed to an arithmetic scale from a log-transformed value) was assumed to be normally distributed with a mean and standard deviation equal to those of the real data. Then we calculated the product of those 10 probabilities and used the joint probability as a summary statistic. Simulated parameter sets were accepted in proportion to the joint probability. For example, if the joint probability was 0.8 for a simulation, the parameter set of the simulation has an 80% chance of being accepted. Because the joint probability can be considered a direct likelihood, this method is superior to the standard ABCs, which depend on the choice of summary statistics and arbitrary tolerance (discussed in Kutsukake & Innan 2013, 2014).

In total, 500 parameter sets were collected for estimating posterior distributions. All results were visualised using R version 3.5.3 (R Core Team 2017). The simulation code was written in C and is available in a public repository (https://github.com/YukiHaba/ABC-PCM).

## Results

### *Model 1:* Logistic functions

When the latent evolutionary function was logistic, the social system of the MRCA was estimated to be solitary in 61.4% of the cases (Fig. 3a, node A). The accepted functions of the transition probabilities varied widely (Fig. 4a), but showed several consistent patterns. First, the curvature parameter of functions, *a*, was positive in all cases (Table 3, Fig. 4b; see Supplementary material for a low risk of type I error), indicating that the probability of transition from a solitary to social species increased as group size increased. In addition, *p*(*x*) typically increased rapidly around *x* = 2 to 4, which corresponded to the peak of the posterior distribution of *b* (Table 3, Fig. 4c).

At each internal node before reaching the common ancestor of social species, the inferred social system was predominantly solitary (Fig. 3a). The common ancestor of solitary species was likely to be solitary (Fig. 3a, node C), and a similar pattern was evident at the node of the common ancestor of *B. janetta* and *B. suillus*. By contrast, at the nodes leading to the clade of *Fukomis*, predominantly social states were inferred (Fig. 3a, node B). The estimated group sizes were consistent with the inferred social systems at internal nodes (Fig. 3b–d); the more social the system was likely to be, the larger the group size.

We detected marginal directional selection for increasing group size in the branch that led to naked mole rats (the proportion of *k*_*n*_ < 1 was 6.4%, Fig. 3f). By contrast, we did not detect significant directional selection in the branch leading to another eusocial species, Damaraland mole rats (the proportion of *k*_*d*_ < 1 was 15.8%, Fig. 3g). Based upon this non-significant result for *k*_*d*_, we repeated the estimation after excluding *k*_*d*_ (i.e., directional selection was assumed only for the branch leading to naked mole rats), but the results did not change qualitatively (data not shown).

### *Model 2:* Exponential functions

With the exponential latent evolutionary functions, the MRCA was inferred to be solitary in 60.4% of the cases (Fig. 5a, node A), similar to model 1. Again, although the accepted functions of the transition probabilities between social and solitary varied (Fig. 6a), the curvature of *p*(*x*), *c*, was always positive (Table 3, Fig. 6b; see Supplementary material for a low risk of type I error).

**Fig. 3.**
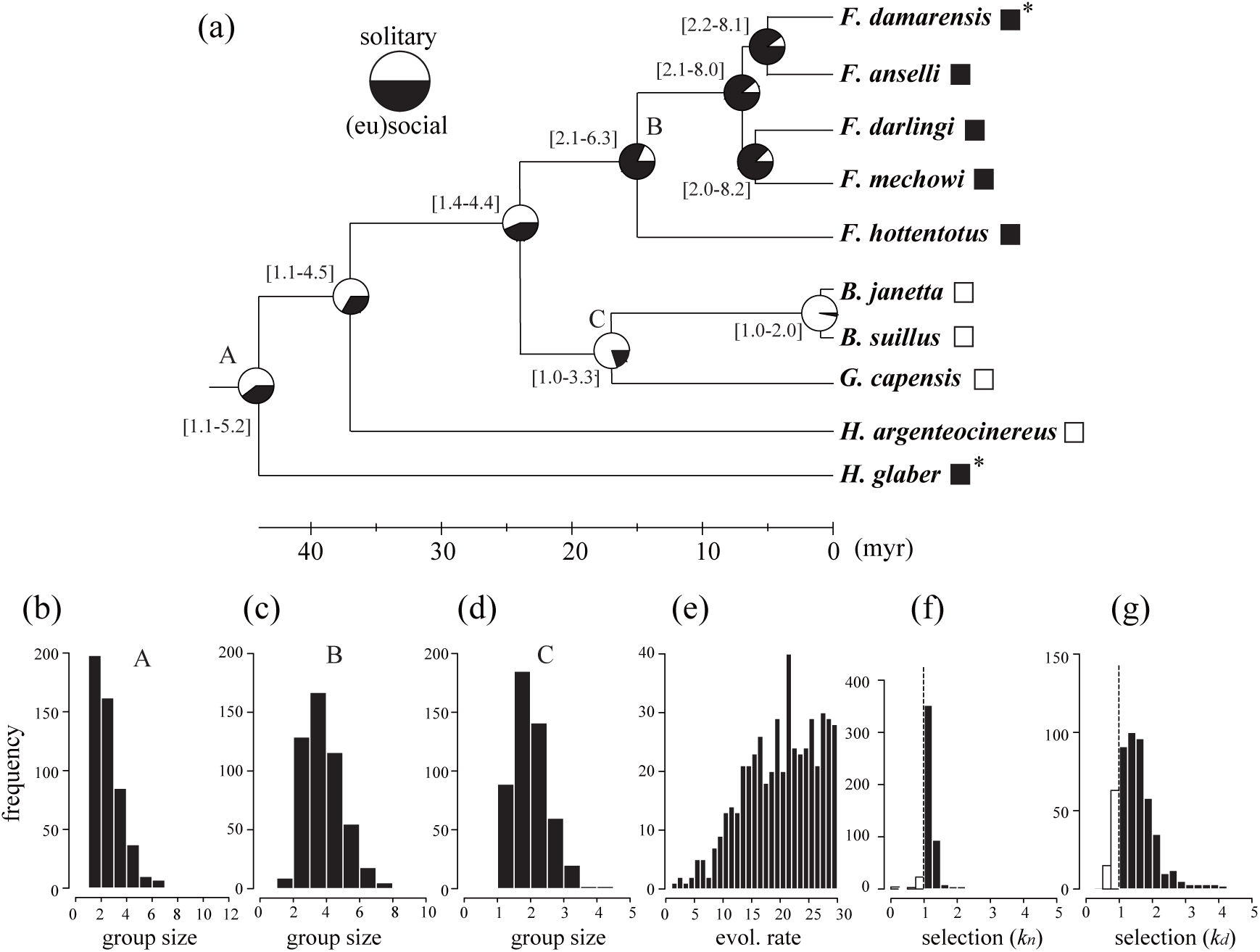
The estimated evolutionary trajectory with the latent function being logistic (model 1). (a) The estimated parameters on the phylogeny. The social state of each tip species is shown as black (solitary) or white (social) squares. Asterisks indicate eusocial species. The proportion of solitary/social states and the 95% CI of the posterior distribution of group size (within brackets) are shown at each internal node. (b–d) The distributions of simulated group size at nodes A (MRCA), B, and C in the phylogeny. (e) The posterior distribution of the baseline evolutionary rate. (f, g) The posterior distributions of the selection coefficients *k*_*n*_ and *k*_*d*_. The dashed line at *k* = 1 indicates the proportion of simulations in which directional selection for larger group size was detected (black histograms).

**Fig. 4.**
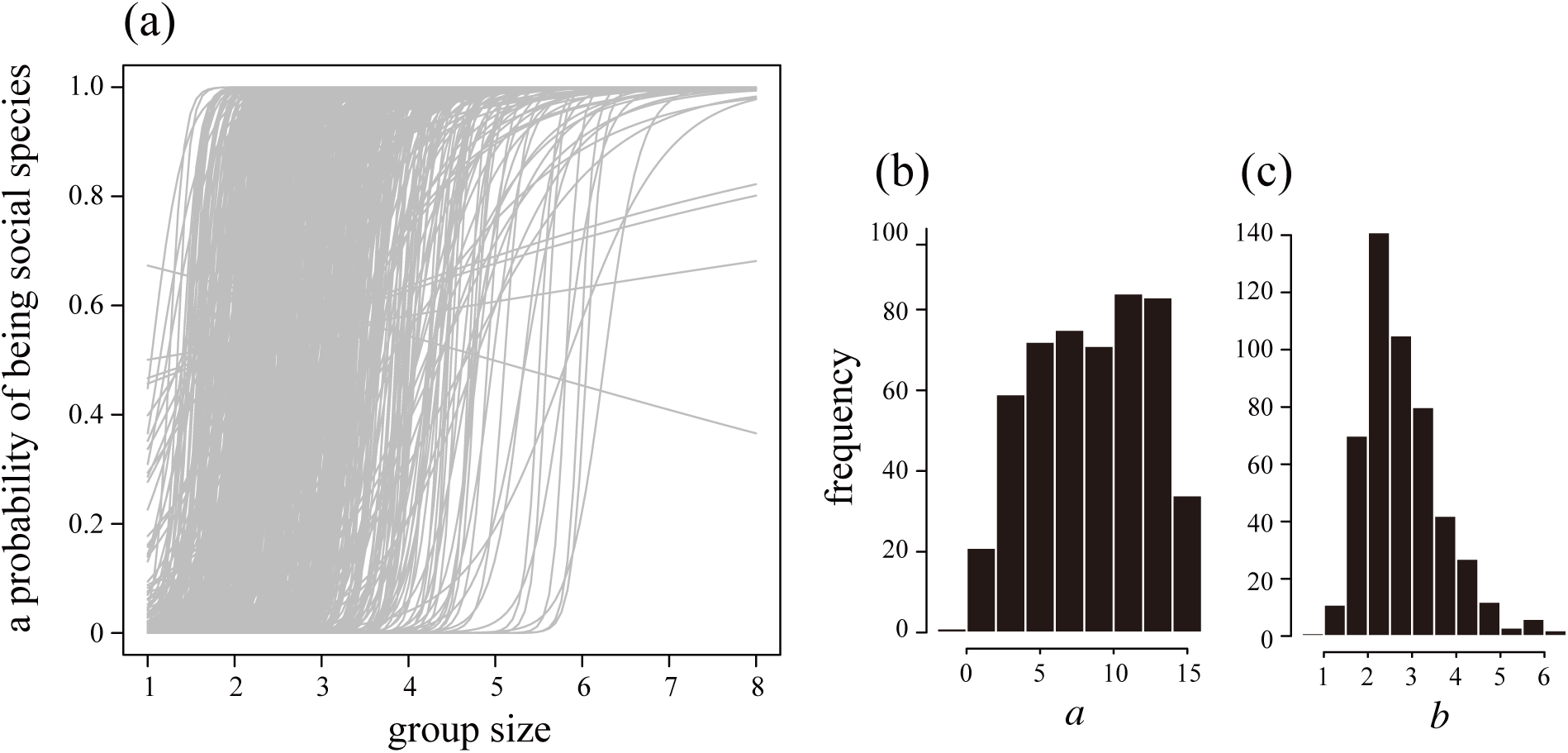
Accepted latent functions in model 1. (a) All 500 accepted latent functions *p*(*x*), i.e., the probability of transition from a solitary state to a social state as a function of group size. Most accepted functions increased steeply at group sizes of 2 to 4. (b, c) The posterior distributions of *a* and *b*, the curvature of the function and the group size at which *p*(*x*) equals 0.5.

**Fig. 5.**
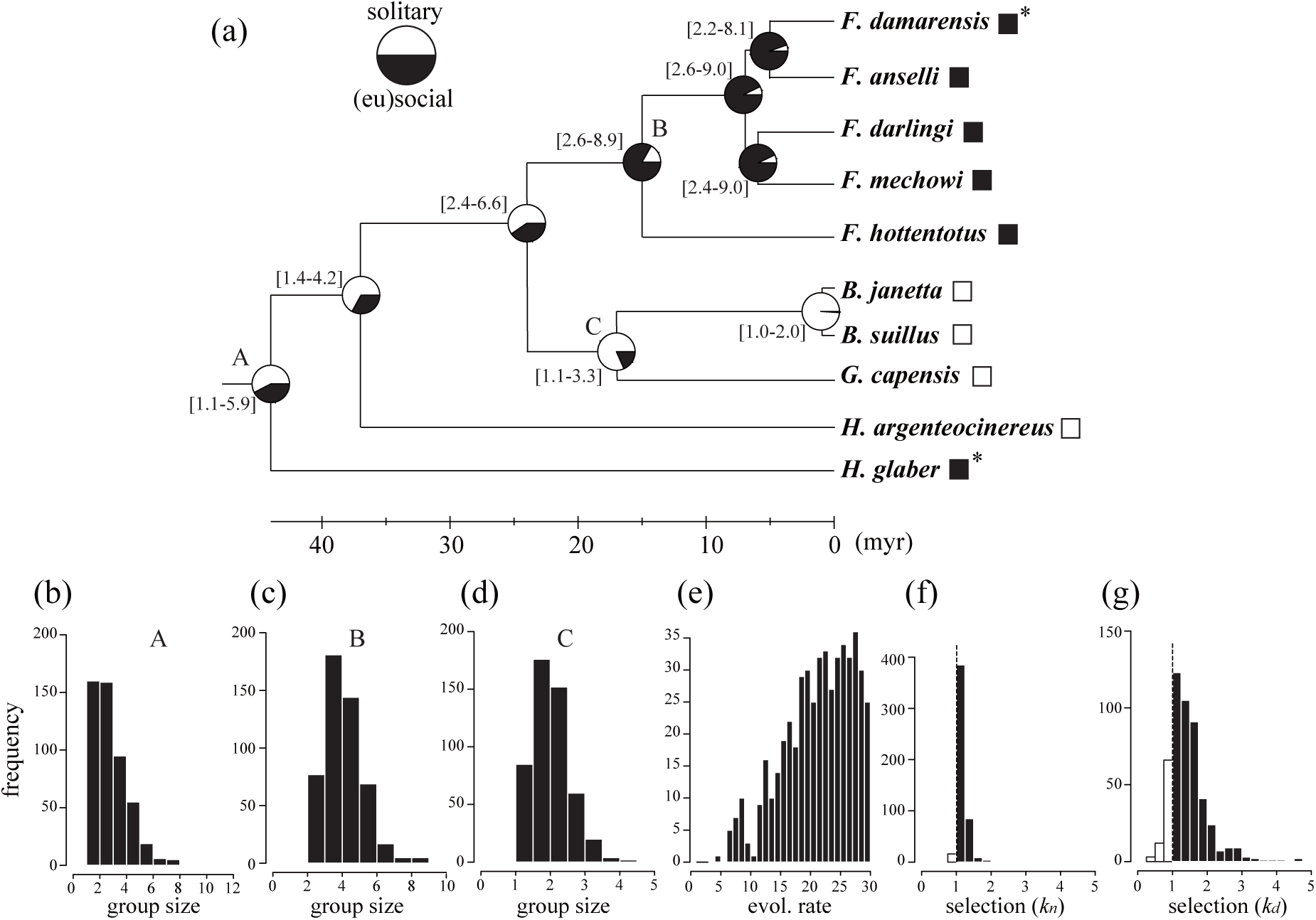
The estimated evolutionary trajectory with the latent function being exponential (model 2). (a) The estimated parameters on the phylogeny. The social state of each tip species is shown as black (solitary) or white (social) squares. Asterisks indicate eusocial species. The proportion of solitary/social states and the 95% CI of the posterior distribution of group size (within brackets) are shown at each internal node. (b–d) The distributions of simulated group size at nodes A (MRCA), B, and C in the phylogeny. (e) The posterior distribution of baseline evolutionary rate. (f, g) The posterior distributions of the selection coefficients *k*_*n*_ and *k*_*d*_. The dashed line at *k* = 1 indicates the proportion of simulations in which directional selection for larger group size was detected (black histograms).

**Fig. 6.**
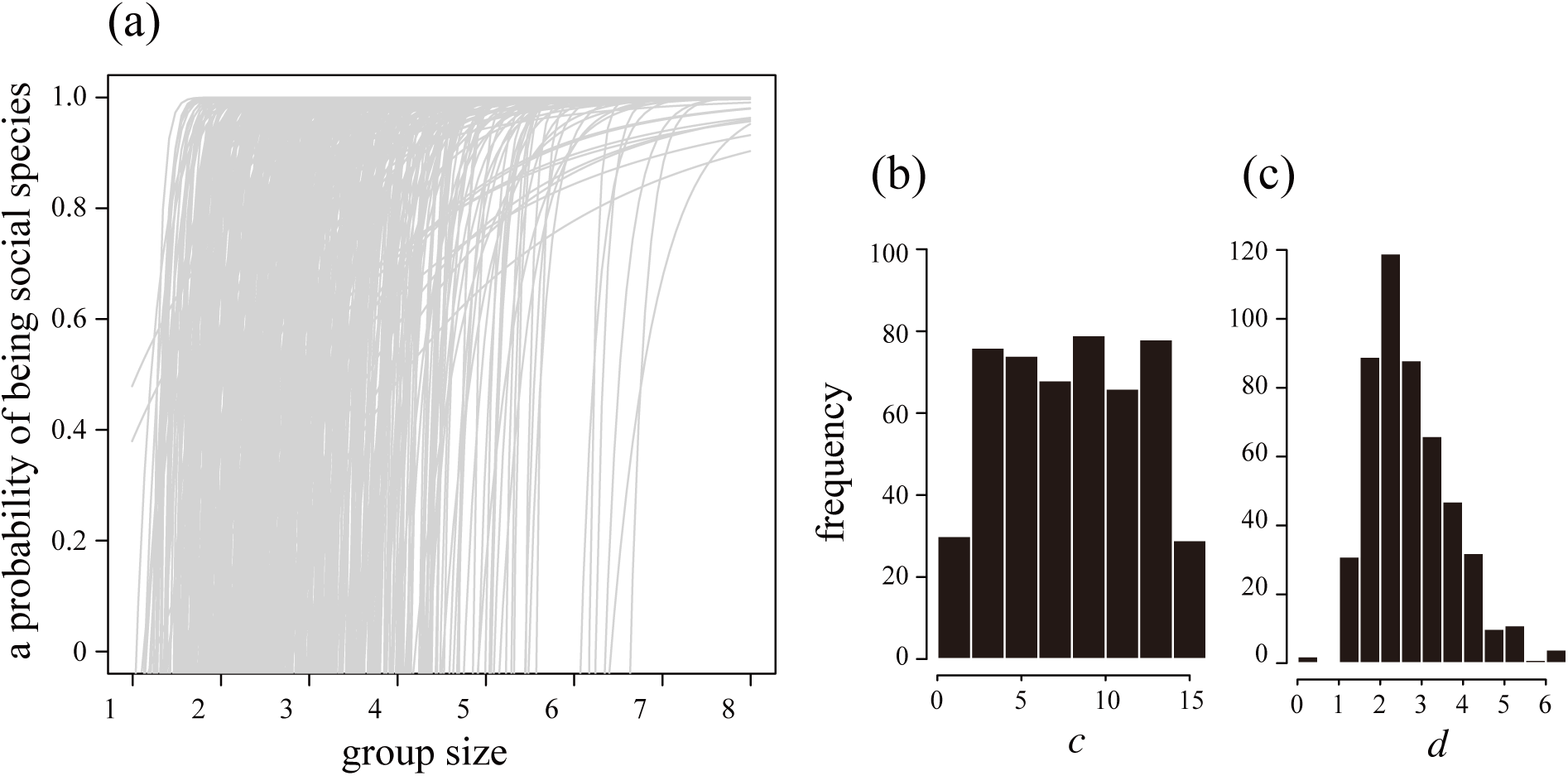
Accepted latent functions in model 2. (a) All 500 accepted latent functions corresponding to *p*(*x*). Similar to the case in model 1, most of the accepted functions increased at a group size of 2 to 4. (b, c) The posterior distributions of *c* and *d*, the curvature of the function and the group size at which *p*(*x*) equals 0.

Group size and sociality at each internal node were almost identical to those in model 1. Within the clade of *Fukomis*, we inferred predominantly social states (Fig. 5a, node B) and relatively large group sizes (Fig. 5c). Within the clade of *Bathyergus* and *Georychus*, in contrast, the social system at each node was consistently solitary (Fig. 5a, node C) and the group size was around 2 to 4 (Fig. 5d).

We detected significant directional selection for increasing group size in the branch leading to naked mole rats (the proportion of *k*_*n*_ < 1 was 3.4%; Fig. 5f). However, we did not detect significant directional selection in the branch to eusocial Damaraland mole rats (the proportion of *k*_*d*_ < 1 was 16.2%; Fig. 5g). Again, based on this non-significant result for *k*_*d*_, we repeated the estimation after excluding *k*_*d*_ (i.e., directional selection was assumed only for the branch leading to naked mole rats); however, the result did not change qualitatively (data not shown).

## Discussion

This study proposes an extension of a previously described ABC-PCM (Kutsukake & Innan 2013, 2014) to analyze complex evolutionary scenarios in which discrete and continuous traits are biologically interwoven. This is the first ABC-PCM framework that allows simultaneous analyses of the interdependent evolution of discrete and continuous traits. In addition, our model offers at least two advantages over other existing methods.

First, this study incorporated a feature that has rarely been included in a PCM framework: branch-specific directional selection (Kutsukake & Innan 2013, 2014; Harano & Kutsukake 2018). This inclusion was possible thanks to the flexibility of the ABC-PCM, which does not require the mathematical expression or analytic solution of a likelihood function. Second, our multivariate model can incorporate flexible user-defined functions that describe the evolutionary relationship between continuous and categorical traits. Previous methods can incorporate nonlinear functions such as logit and probit functions (Ives and Garland 2010; Hadfield & Nakagawa 2009; Hadfield 2015), but the choice of a latent function is less flexible than in our present method. Another well-established method for simultaneously analyzing both continuous and categorical traits is the threshold model (Felsenstein 2005, 2012). The threshold model is equivalent to a phylogenetically controlled linear model with a probit latent function (see Supplementary Material in Hadfield 2015 for a useful graphical representation). Although the assumption is mathematically convenient for analyses, it may be an oversimplification of the evolutionary relationship between the continuous and categorical traits of interest.

Applying this new approach to data on African subterranean rodents, we estimated the intricate evolutionary trajectory of sociality and group size. Two models with different latent evolutionary functions were tested: logistic and exponential. Although the exponential function has not been used in previous PCMs, we believe that this function is suitable for our study species for two reasons. First, group size should have a minimum value of one, and second, a species whose group size is one must be a solitary species. Although these functions have different mathematical characteristics and are not nested, they have common features such as a monotonic increase and an asymptotical approach from one to zero. Potentially due to these common features, the estimated parameters showed similar posterior distributions. In both models the social state of the MRCA was not decisively solitary or social, and its estimated group size varied (node A in Figs. 3a and 5a).

The latent evolutionary functions *p*(*x*) and *q*(*x*) had a consistent pattern in both models (Figs. 4 and 6). Namely, most of the accepted functions had a steep transition around a group size of 2 to 4. This indicates that this particular window of group size was an evolutionary tipping point of the transition between solitary and social states. This study is the first to quantitatively infer the range of group size that is crucial to the evolution of sociality in this clade.

We tested differential selective pressure on group size in the branches leading to the two eusocial species in this clade (*k*_*n*_ and *k*_*d*_); we detected directional selection (*k*_*n*_) for larger group size in the branch leading to naked mole rats in model 2 (Fig. 5f) but not in model 1 (Fig. 3f). In both models, a selective pressure was not detected in the branch leading to Damaraland mole rats (*k*_*d*_, Figs. 3g and 5g). Thus, although both species are deemed eusocial, the two species may have undergone qualitatively different evolutionary paths (Burda *et al*. 2000). Future studies should explore the differences in the ecological and evolutionary causes of the evolution of these two eusocial species.

One critical limitation of this case study was the relatively low sample size (10 species; see Supplementary material). This sample size is caused by three inevitable limitations. First, the family Bathyergidae is a monotypic group composed of less than 20 OTUs, which is an inevitable constraint on sample size. Second, detailed data for group size are not available for all species of this family, which further limits the available data. Finally, and most importantly, expanding the number of study species to include closely related non-subterranean species is questionable, as it is highly likely that the underground ecological niche has affected predation pressure and consequently sociality. More broadly, it remains unclear whether heterogeneity of traits other than the traits of interest would affect biological interpretation in PCM studies based on a large sample size. It is generally believed that larger sample sizes (a larger number of species, or more precisely, a larger number of evolutionary transitions) would produce a more powerful test in PCMs. While this is true, it remains largely undiscussed how the inclusion of species that have fundamentally different ecological or behavioral traits would affect the results. In this sense, in addition to a large-scale comparison, it is important to conduct PCMs that focus on a taxon that shares fundamental traits, even if the sample size is not very large.

One major challenge in ABC-PCMs is that they are computationally intensive. In our study, for example, it took several days to accept one parameter set (MacPro, OS 10.6.7, 2 *x* 2.93 GHz Quad-Core Intel Xeon; also see Kutsukake & Innan 2013). To compare the performance of our model to an available method, we ran a bivariate model on our dataset and phylogeny using MCMCglmm (Hadfield & Nakagawa 2009; Hadfield 2015) assuming that group size and sociality follow Gaussian and probit (“threshold”) families, respectively. For simplicity, we ran the program without incorporating intraspecific variation, branch-specific selection, or the bounding of group size. The MCMCglmm results qualitatively agreed with our results (e.g., 95% CI of the covariance between group size and sociality > 0), yet the process took only a few minutes to an hour on a standard laptop, depending on the parameters (results not shown). Thus, the scope of the present ABC-PCM algorithm is currently limited to analyses of relatively small to moderate numbers of species (29 species in Harano & Kutsukake 2018), but it is not suitable for a large number of species. Improvements in both efficient simulation algorithms and computational power will enable analyses based on a larger number of species.

In summary, we have developed a flexible multivariate ABC-PCM that has great potential for testing biologically intricate scenarios of trait evolution. Although our analyses considered the evolutionary transition of sociality in a relatively small number of species, our method can be applied to other topics and to larger datasets. Despite the fact that our analyses focused on a simple case in which a continuous trait affects the state of a discrete trait, other causational patterns can be dealt with by a flexible setting of an evolutionary simulation. Furthermore, our ABC-PCM framework can also be extended to model more complex evolutionary trajectories, such as asymmetric transitions between states and/or more than two states of discrete traits with different transition functions. Thus, our method allows evolutionary biologists to explore various hypotheses of interest concerning the evolution of interrelated traits.

## Acknowledgements

We would like to thank Dustin R. Rubenstein, Rafael Maia, and Margaret E. O’Brien at Columbia University and Jessica Zung at Princeton University for useful comments and advice. Masahito Tsuboi at University of Oslo also provided useful insights on the earlier version of manuscript. Two reviewers gave detailed and helpful comments on the manuscript. This study was financially supported by MEXT (No. 25711025) to K. N.

## Author contributions

Y. H. and K. N. conceived, designed, and performed the analysis. Both discussed the result and wrote the final manuscript.

