## Supplementary Material for "A multivariate phylogenetic comparative method incorporating a flexible function between discrete and continuous traits"

**for**

### Power analyses

We examined the statistical performance of the proposed ABC-PCM by conducting power analyses for a situation that resembled the analyzed data. Specifically, we were interested in whether this method could successfully estimate $a$ (model 1) and $c$ (model 2). Below, we assumed that the discrete trait has two states, $A$ and $B$. The phylogeny used in these power analyses were the same as that for the analyzed data.

First, we determined a parameter set that included the MRCA ($\theta_0$), evolutionary rate ($\mu$), and parameters of the latent function ($a$ and $b$ for model 1; $c$ and $d$ for model 2). We fixed the value of the MRCA ($\theta_0$) to 0, evolutionary rate ($\mu$) to 10, and $b$ and $d$ to 1. Because we were interested in how parameters $a$ and $c$ would affect the performance of our method, we changed these parameters to values ranging from 0.0001 to 100 (see Fig. S1A, C). We selected the value range from that of an approximately independent relationship to the pattern similar to the threshold model; small values of $a$ or $c$ approximately correspond to a biological situation in which the discrete and continuous traits evolved independently (Fig. S1A, C). Sufficiently large values of $a$ or $c$ are close to a model where the discrete trait is determined by a single value of the continuous trait (Fig. S1A, C). The power analyses did not include the directional selection to avoid excessive complexity. For simplicity, we did not set a boundary condition for the evolutionary range of trait values (i.e., the continuous trait value can be negative).

We conducted 600 rounds of ABC estimation. In each round, we first made one “target” dataset by a trait simulation using the parameter set pre-determined by ourselves. As a result, we obtained the target dataset with both discrete and continuous variables whose inter-relationships were determined by the latent function used in model 1 or 2. Using the “target” data, we conducted ABC-PCM analyses and estimated a posterior distribution for each parameter by accepting 200 parameter sets.

This procedure was subject to practical complications. One was that the target data differed across rounds, by which the number of species with two discrete traits ($A$ and $B$) also varied. In our preliminary survey, we noticed that the statistical performance was strongly affected by the number of tips with different discrete categories. Therefore, we calculated the power separately according to the distribution of discrete categories at the tips. Second, we also found that the tolerance for judging the acceptance or rejection differed across rounds because of differences in the target data (Fig. 2, Step 4). To address this problem, we first conducted a preliminary ABC estimation for one million rounds and employed the best summary statistics as the tolerance for the continuous trait. We required a complete match between the target and generated data for the discrete trait as in the main analyses.

After accepting the 200 parameters for each round, we checked whether the 95% credible interval of a posterior distribution overlapped with zero. We repeated this procedure (i.e., generate data, ABC, check the 95% CI) 600 times and calculated the proportion for which the 95% credible interval exceeded zero. We reported the powers calculated from the dataset with four species of state A (which corresponds to solitary species in analyses of real data) and six species of state B (which corresponds to social species).

As a result, our ABC had a low proportion (<0.05) of detecting a positive value of $a$ and $c$ when a relationship between discrete and continuous traits were almost independent (e.g., $a = 0.001$ and 0.1 and $c = 0.01$; see the shaded area in Fig. S1 B and D). This indicates that this ABC does not have a problem with type I error. By contrast, the proportions of detecting positive $a$ and $c$ (i.e., power) were approximately 0.140 (for model 1) and 0.163 (for model 2) within the range of the estimated CI for $a$ and $c$ (see Table 3 in the main text).

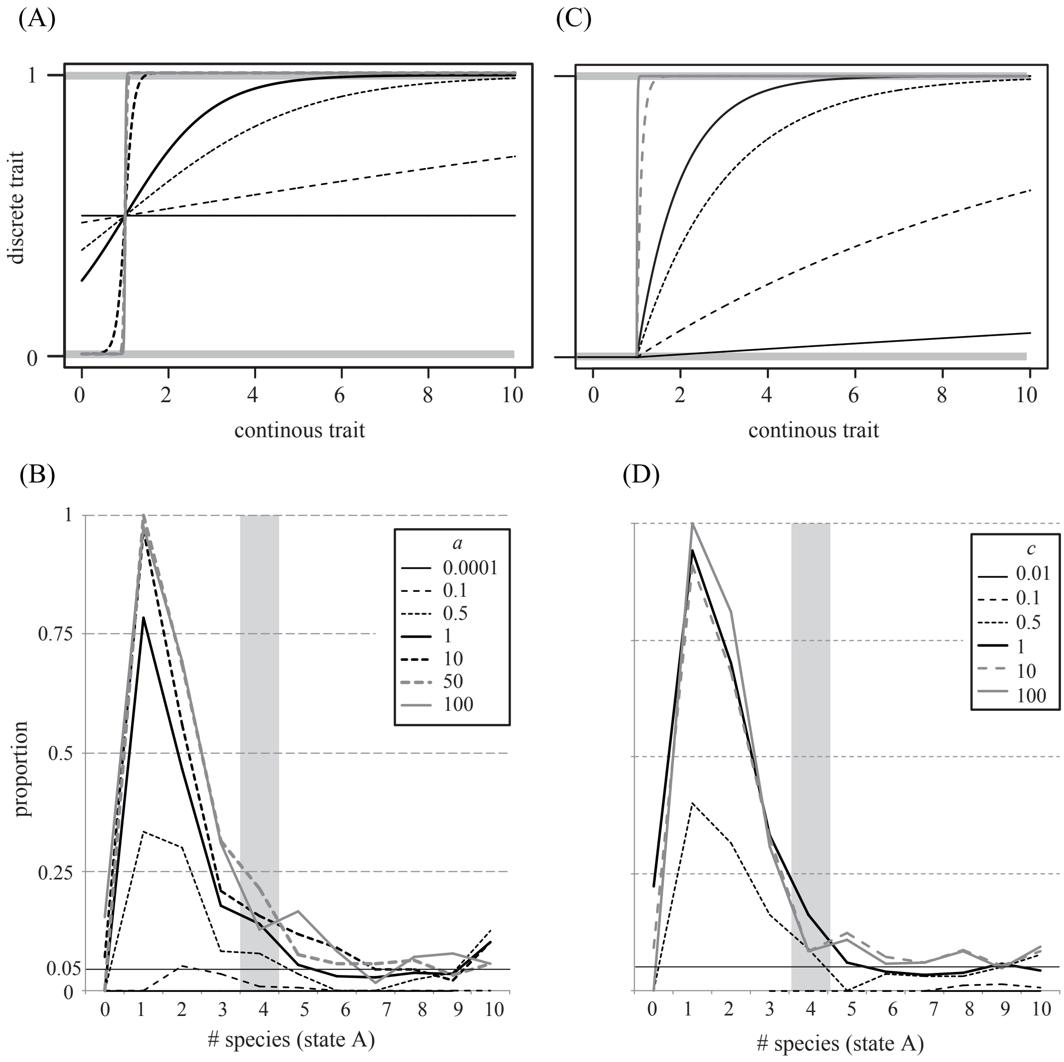


**Fig. S1.** The power of the proposed PCM increased as the latent function became steeper. The proportion of ABC estimation in which the 95% CI of *a* or *c* did not overlap with zero (vertical axis) is shown separately according to the number of species with two discrete traits (on the horizontal axis) and the pre-determined value of *a* or *c* (indicated by different lines). The shaded area corresponds to the cases in which the generated data have four and six states of a discrete variable, similar to our data analyses of social evolution in African mole rats. Prior distributions were set as U(-20, 20) for MRCA, U(0, 20) for evolutionary rate, and U(-10, 10) for *b* or *d*. (A) Shapes of logistic functions. (B) Results of model 1 (logistic function). For parameter *a*, we used the following prior distribution. U(-10, 10) for *a* = 0.0001, 0.1. 0.5, 1; U(-50, 50) for *a* = 10; U(-200, 200) for *a* = 50; and U(-300, 300) for *a* = 100. (C) Shapes of exponential functions. (D) Results of model 2 (exponential function). For the parameter *c*, we used the following prior distribution: U(-10, 10) for *c* = 0.01, 0.1. 0.5, 1; U(-50, 50) for *c*=10; and U(-300, 300) for *c* = 100. Note that the power could not be calculated for some combinations of the discrete trait because of too rare acceptance.
